# Bioproduction of single-stranded DNA from isogenic miniphage

**DOI:** 10.1101/521443

**Authors:** Tyson R. Shepherd, Rebecca R. Du, Hellen Huang, Eike-Christian Wamhoff, Mark Bathe

## Abstract

Scalable production of gene-length single-stranded DNA (ssDNA) with sequence control has applications in homology directed repair templating, gene synthesis and sequencing, scaffolded DNA origami, and archival DNA memory storage. Biological production of circular single-stranded DNA (cssDNA) using bacteriophage M13 addresses these needs at low cost. A primary goal toward this end is to minimize the essential protein coding regions of the produced, exported sequence while maintaining its infectivity and production purity, with engineered regions of sequence control. Synthetic miniphage constitutes an ideal platform for bacterial production of isogenic cssDNA, using inserts of custom sequence and size to attain this goal, offering an inexpensive resource at milligram and higher synthesis scales. Here, we show that the *Escherichia coli* (*E. coli*) helper strain M13cp combined with a miniphage genome carrying only an f1 origin and a β-lactamase-encoding (*bla*) antibiotic resistance gene enables the production of pure cssDNA with a minimum sequence genomic length of 1,676 nt directly from bacteria, without the need for additional purification from contaminating dsDNA, genomic DNA, or fragmented DNAs. Low-cost scalability of isogenic, custom-length cssDNA is also demonstrated for a sequence of 2,520 nt using a commercial bioreactor. We apply this system to generate cssDNA for the programmed self-assembly of wireframe DNA origami objects with exonuclease-resistant, custom-designed circular scaffolds that are purified with low endotoxin levels (<5 E.U./ml) for therapeutic applications. We also encode digital information that is stored on the genome with application to write-once, read-many archival data storage.

## Introduction

Kilobase-length, single-stranded DNA (ssDNA) is essential to numerous biotechnological applications including sequencing (1), cloning (2), homology directed repair templating for gene editing (3), DNA-based digital information storage (4,5), and scaffolded DNA origami (6–9). Specifically, scaffolded DNA origami enables the fabrication of custom structured nanoscale objects with application to nanoscale lithography (10,11), light harvesting and nanoscale energy transport (12–15), metal nanoparticle casting (16), and therapeutic delivery (17,18). In this approach, a long ssDNA scaffold is folded via self-assembly into arbitrary, user-specified shapes by slow annealing in the presence of complementary short oligonucleotide “staples”. These staples are designed using Watson-Crick base-pair complementarity to the scaffold, forcing sequences that are far apart in sequence space to be close in physical space. Fully automated, top-down computational design of scaffolded DNA origami nanostructures has now been enabled by sequence design algorithms in both 2D and 3D (19–23), enabling the democratization of otherwise complex scaffolded DNA origami design that previously excluded non-experts. However, therapeutic and materials science applications that require large-scale, low-cost scaffold with custom length and sequence requirements are still hindered by limitations in production of circular, isogenic scaffolds on the 1–3 kb scale (24–26).

The most common low-cost source of native cssDNA for scaffolded DNA origami is the 7,249 base single-stranded M13mp18 phage genome, which has allowed production of up to 410 mg of cssDNA per liter of *E. coli* growth through fed-batch fermentation (27). Innovative approaches to achieving custom bacterial scaffolds (25) and staple production were recently implemented (26). However, numerous applications require increased purity of circular custom scaffolds, such as in therapeutic applications of DNA origami, that would additionally require synthetic staples for chemical functionalization and stabilization (28). Further, elimination of the antibiotic selection marker from the produced linear (26) or circular (25) ssDNA does not allow for subsequent re-infection without this selective control (25), potentially limiting downstream biological applications where reinfection would be useful, such as for biological sequence amplification.

These preceding advances employed helper plasmid systems where the M13 coding sequences are sub-cloned onto a double-stranded, low-copy number vector that is co-transformed with a phagemid containing an ssDNA origin of replication (e.g., f1 origin) that allows for the synthesis and packaging of ssDNA. The most commonly used helper plasmid system is M13KO7 (29), which maintains a packaging sequence, albeit with a mutated packaging signal to reduce the packaging frequency. This system has shown utility in phage display (30–32), and has also been applied to produce a phagemid that encodes a 2,404-nt sequence containing an cssDNA f1 origin, a dsDNA pUC origin, and an ampicillin selection marker (24). However, the preceding phagemid cssDNA was contaminated by other DNAs, both from the dsDNA phagemid and M13KO7 (24). Importantly, isolation of the target ssDNA from these DNA impurities would require subsequent purification steps for further bioproduction scale-up, and would potentially introduce high background in sequencing application, with possible errors in phage propagation, and lower yields in DNA origami applications. To overcome these potential problems with the helper system, the ssDNA origin of replication and packaging signal was entirely removed from a helper plasmid (*E. coli* str. M13cp) (33), thereby enabling the biological production of isogenic cssDNA without DNA impurities. Indeed, recent advances in engineered phage systems have utilized this strategy (25,26), however, purification would ultimately be required for DNAzyme-based approaches to bacterial scaffold as well as staple production (26) and optimization for pure cssDNA production (34) without genomic or plasmid contamination is still required (25,26). Thus, there remains a critical need for scalable production of isogenic cssDNA at the 1–3 kb length scale that maintains replication capacity.

To address this need, in the present work we show that isogenic miniphage production of cssDNA is scalable by fermentation using the *E. coli* str. SS320 with the M13cp helper plasmid (generous gift of Dr. Andrew Bradbury, Los Alamos National Lab). Three miniphages were synthesized using both classic restriction and restriction-free (RF) cloning (35,36). The miniphage presented here maintain the selection marker and origin of replication, which allows for the reinfection of the phage in culture while reducing the occurrence of contaminating dsDNA because they do not contain a double-strand origin of replication, similar to the natural M13 phage. Monitoring phage yields and growth rates of the bacteria, we identify an 8-hour timepoint after inoculation that yields maximal cssDNA production with no detectible DNA contamination. Silica-column-based DNA extraction techniques from clarified media yielded 2 mg of pure cssDNA per liter of culture with < 5 endotoxin units per milliliter of sample. We demonstrate that the cssDNA material is practical for the generation of custom length circular scaffolds for wireframe DNA origami with partial sequence control, with additional applications to write-once, read-many archival DNA data storage.

## Results

A variant of extension-overlap, restriction-free cloning (35,36) using long ssDNA (**Figure 1a**) was applied for the *de novo* assembly of a miniphage genome containing only an f1 origin of replication and an ampicillin resistance selection marker (phPB52). Two kilobase-scale megaprimer ssDNAs were generated using asymmetric PCR (aPCR; (37)) using 5′-phosphorylated primers: a top-strand megaprimer encoding the f1 origin sequence (427 nt; (38)) and a bottom-strand megaprimer encoding an ampicillin resistance cistron (*bla*; 1,249 nt) (**Figure 1b**, **Supplementary Figure S1**). The kilobase primers were synthesized such that the two sequences contained a complementary sequence of 20 nt on each of the 5′ and 3′ ends (**Figure 1a**). The two megaprimers were mixed at equimolar concentration and completed to dsDNA using PCR, followed by enzymatic ligation. The ligated plasmid (**Supplementary Figure S2**) was transformed into chemically competent *E. coli* str. M13cp (33) and dual selected on ampicillin and chloramphenicol with no detectable toxicity due to the miniphage, resulting in normal colony shape and size. Two out of eight colonies screened were found to be of the exact sequence desired. Liquid culture was inoculated and grown in a shaker flask, after which the culture was centrifuged to separate the phage-containing media from the bacterial pellet. Phage in the clarified media were visualized by TEM, showing the anticipated size of and homogeneity (**Figure 1c and Supplementary Figure S1**). The cssDNA from this phage was isolated using silica column purification and showed 88% cssDNA purity according to agarose gel imaging (**Figure 1d**), with an approximate yield of 0.5 mg per liter of bacterial growth, while the bacterial pellet showed helper plasmid, dsDNA intermediate phage DNA, and cssDNA.

**Figure 1.**
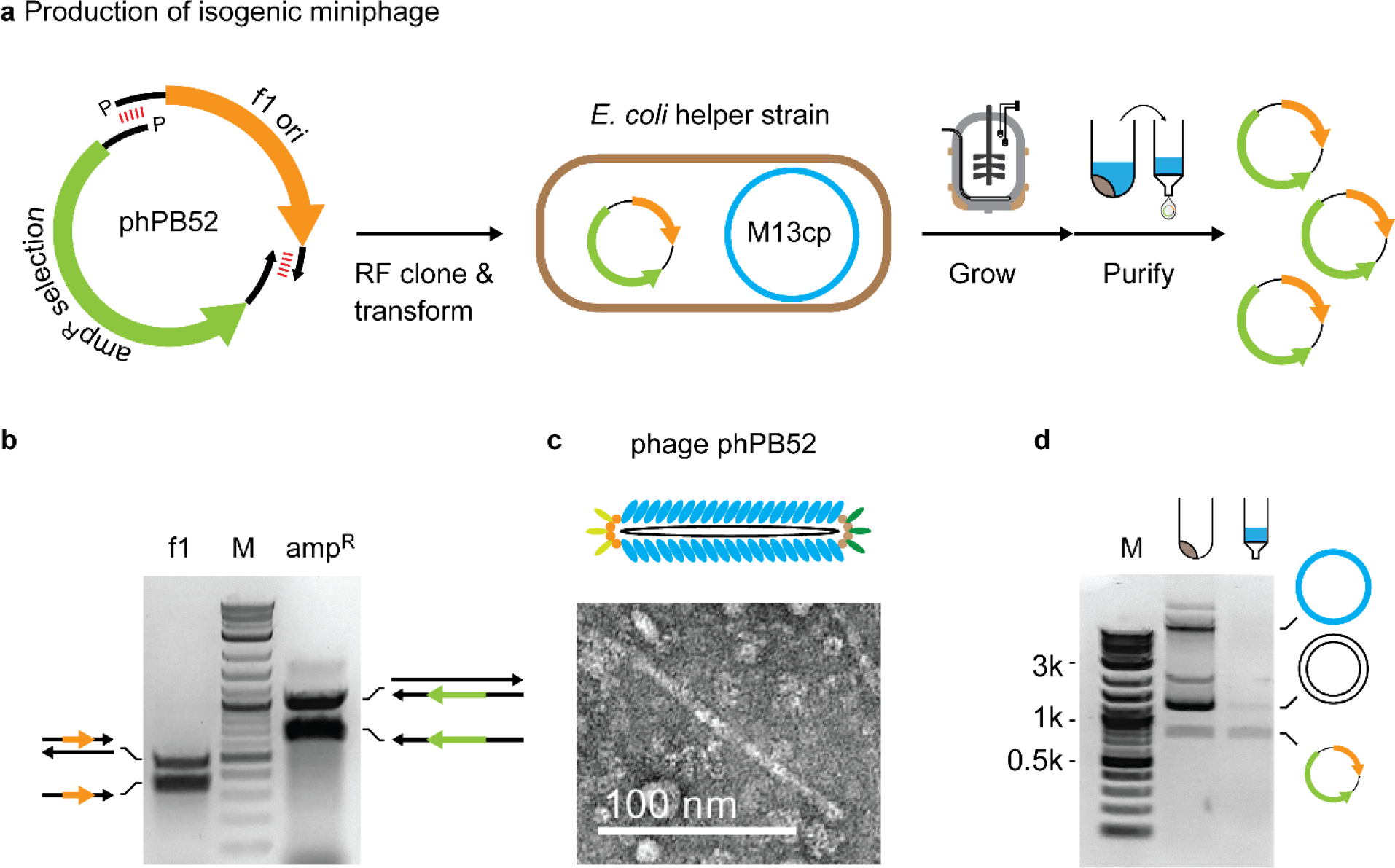
Scalable bacteriophage production of isogenic cssDNA. (**a**) Miniphage phPB52 was assembled using restriction-free (RF) cloning was used for miniphage phPB52 assembly and transformed into *E. coli* containing the M13cp helper plasmid for production of isogenic cssDNA. (**b**) aPCR was used to generate the two ssDNA megaprimers for RF cloning encoding the f1 origin of replication (f1 ori) and the *bla* ampicillin selection marker (Selection). (**c**) Phage particles from clarified media were visualized by TEM. (**d**) DNA purification from the bacterial pellet and the clarified media show mostly pure cssDNA in the media and cssDNA and dsDNA phagemid, and helper plasmid in the bacterial pellet. See **Supplementary Figure S1**for uncropped images.

Having generated a phage containing only the f1 origin and a resistance gene, demonstrated to be stably produced and exported to the media from the helper strain, we next sought to generate a second phage with a synthetic fragment of DNA that is orthogonal in sequence to bacterial and phage genomes. A fragment of length 844 nt was ligated between the f1 origin and the *bla* cistron using standard restriction cloning to generated a plasmid of size 2,520 nt (phPB84; **Figure 2a** and **Supplementary Figure S2**). This plasmid was transformed into the helper strain and the produced phage was purified and its sequence verified by primer walking with Sanger sequencing (**External Table S1** and **S2**)

**Figure 2.**
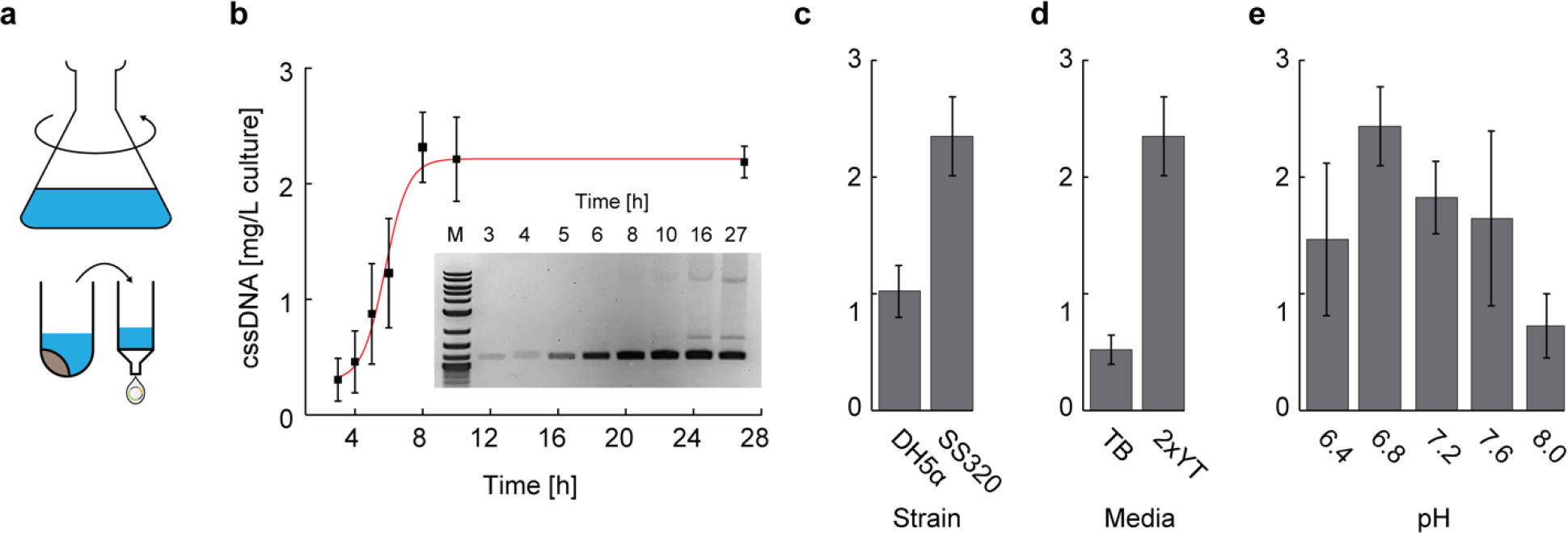
Shaker flask production of pure cssDNA. (**a**) Shaker flask growth of phPB84 was used to optimize conditions for phage amounts and purity. (**b**) Time-course assay of cssDNA production of phPB84, with cssDNA yield calculated by absorbance at 280 nm and purity adjusted by agarose gel band intensity, showing maximum yield and purity at the 8-hour timepoint. The 16 h time-point is from a separate culture, and therefore is not included in the plot. (**c**) Comparison between DH5a F′Iq and SS320 showing two-fold yield increases in the SS320 strain. (**d**) Comparison between growth media showing five-fold improved cssDNA yield in 2×YT after 8 hours of production. (**e**) Comparison of five pH values for cssDNA production, controlled by use of 100 mM HEPES-NaOH. Error bars indicate standard deviation of triplicate experiments. See **Supplementary Figures S3** and **S4** for uncropped triplicate measurements of all experiments.

In order to obtain milligram-scale production of cssDNA with high genetic purity of the final material, we used a shaker flask setup (**Figure 1a**) to vary the growth time, the *E. coli* strain, the growth media, and the media pH to determine optimal conditions. We found the highest and purest yield of cssDNA production occurred at the 8-hour timepoint after inoculation, near the end of log phase, with production falling off thereafter and the appearance of dsDNA contaminations in the media visualized at the 12-hour timepoint (**Figure 2b** and **Supplementary Figures S3**). Two strains were tested for production: DH5a F′Iq (Invitrogen) and the SS320 strain (Invitrogen). Both express the F pili and are commonly used for phage production, and each was transformed with the M13cp helper plasmid purified from *E. coli* str. M13cp. Strain SS320 showed approximately double the cssDNA yield (**Figure 2c** and **Supplementary Figure S4**) and was therefore chosen as the strain for further optimization of growth conditions. Terrific broth (TB) and 2× yeast extract tryptone (2×YT) media for bacterial growth and cssDNA production were both evaluated for phage growth while also monitoring dsDNA contamination using agarose gel analysis (**Figure 2d** and **Supplementary Figure S4**). Notably, TB had significant dsDNA contamination by the 8-hour timepoint (**Supplementary Figure S4**), and 2×YT was therefore chosen as the optimal media for batch production. Next, we investigated the pH sensitivity of the production of phage material (34), which exhibited a three-fold increase in yield at pH 6.8 and 7.2 compared to pH 8 (**Figure 2e** and **Supplementary Figure S4**).

Having identified the optimal growth conditions in the shaker flask setup, we next identified conditions for scale-up in a batch fermenter process (**Figure 3a**) using a Stedium Sartorius 5L fermenter (Sartorius, Germany). Shaker flask conditions were transferred to the bioreactor setup including using 2×YT media, while pH 7.0 was controlled using external phosphoric acid and ammonium hydroxide. The growth curve was monitored using O.D.600 absorbance measurements and the pH and dissolved oxygen were monitored by calibrated probes. Each timepoint was additionally monitored for cssDNA and dsDNA production using agarose gel analysis (**Figure 3b** and **Supplementary Figure S5**), showing maximal cssDNA yield at the 8-hour timepoint, as with the shaker flask, with minimal contaminating dsDNA up to the 12-hour timepoint (**Supplementary Figure S5**). Extraction of 900 mL of media for phage purification was carried out at the 8-hour timepoint and processed using a silica-column based approach specifically designed to reduce endotoxin levels (EndoFree Megaprep Kit, Qiagen, MD). Gel band intensity analysis after kit purification showed no detectable dsDNA contamination (**Figure 3c**). Sanger sequencing by primer walking verified the sequence of the phage DNA (**External Table S1** and **S2**). The kit-based purification yielded 2 mg of cssDNA/L of culture, matching the yield from phenol-chloroform extraction. Endotoxins were tested using a colorimetric assay (ToxinSensor Chromogenic LAL Endotoxin Assay Kit, GenScript, NJ), showing the final product yielded endotoxin levels at 1.1 ± 0.1 E.U./ml per cssDNA concentration of 10 nM, similar to endotoxin reduction by Triton-X114 ((39); **Supplementary Figure S6**). Circularization of the produced cssDNA was verified by incubation with exonuclease I, showing no detectable degradation after 30 min (**Figure 3d**).

**Figure 3.**
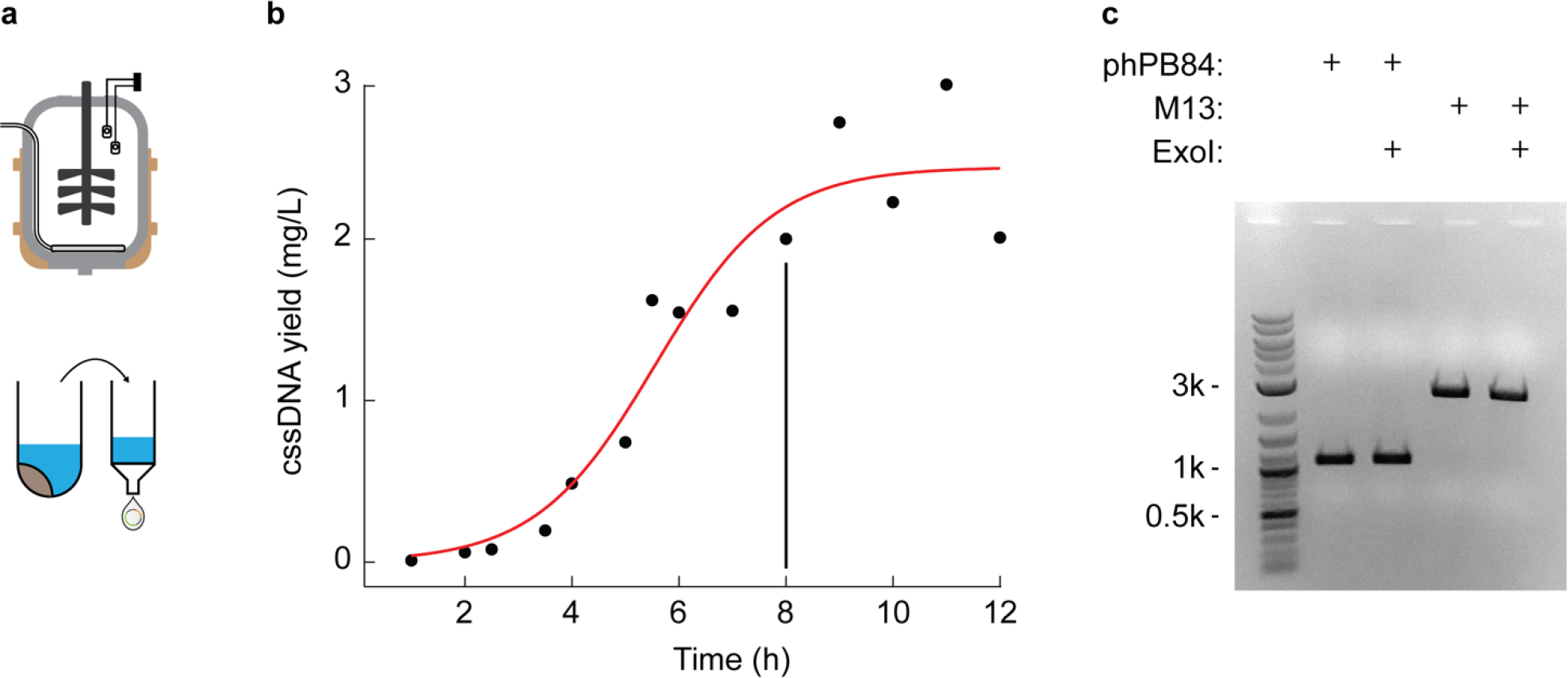
Batch fermenter production of pure cssDNA. (**a**) Scalable production in a stirred-tank bioreactor. (**b**) Time-course assay of cssDNA yield based on agarose gel band intensity analysis (**Supplementary Figure S5**), with the 8-hour timepoint used for 900 mL cssDNA purification. (**c**) Silica-column DNA purification from the PEG-precipitated phPB84 phage showed no detectible dsDNA contamination, similar in purity to commercially available M13mp18, yielding 2 mg of DNA per liter of culture at the 8-hour timepoint. Stability from exonuclease I (ExoI) degradation after 30 min coincubation indicates the ssDNA is circular.

Having implemented a method for milligram-scale production of isogenic miniphage cssDNA, we next applied the method to produce custom length single-stranded DNA scaffold with partial sequence control for application to wireframe scaffolded DNA origami (**Figure 4**). We used the DAEDALUS design algorithm (19) to design a DNA-scaffolded pentagonal bipyramid with a 52-bp edge length (1,580 nt scaffold length) using the smallest phPB52 phage genome sequence (1,676 nt) and a second DNA-scaffolded pentagonal bipyramid with an 84-bp edge length (2,520 nt scaffold length) using the phPB84 phage genome sequence (2,520 nt). Notably, any DNA origami with scaffold lengths larger than 1,676 nt can have perfectly matched phage genome lengths, as exemplified in the pentagonal bipyramid with an 84-bp edge length. DNA origami object folding was characterized using agarose gel mobility shift assays and transmission electron microscopy (TEM), which confirmed monodispersed object sizes with near quantitative yield of self-assembly (**Figure 4a** and **Supplementary Figures S7** and **S8**).

**Figure 4.**
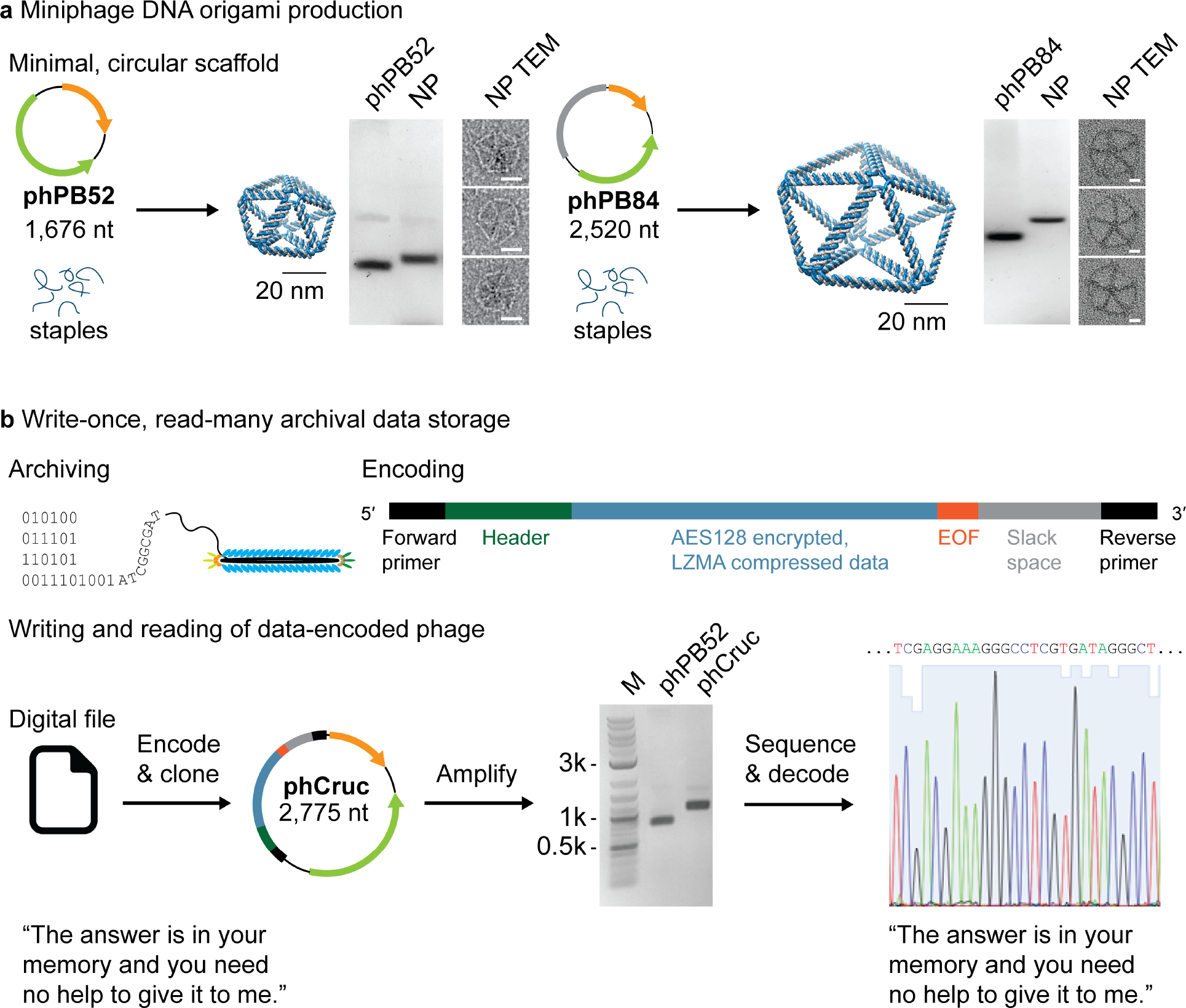
Applications of scalable, isogenic miniphage production. (**a**) Two pentagonal bipyramids of 52-bp and 84-bp edge-lengths were folded using the phPB52 and phPB84 as scaffolds, respectively. Agarose gel shift mobility assays and TEM were used to validate the folding of the scaffold to the expected design. (**b**) Phage particles are natively protected from environmental degradation and easy to amplify by bacterial infection, and thus provide a compelling method for archival and amplification of digital information encoded in DNA. The encoding scheme shown here can generate bio-orthogonal sequences that are designed to limit secondary structure and recombination sites. A digital text file containing a line from <u>The Crucible</u> (40) was ligated to the phPB52 vector and subsequently bacterially produced and amplified. Sanger sequencing by primer walking was used to retrieve the original digital file. See **Supplementary Figures S7** and **S8** for uncropped gel images and example full field TEM micrographs, respectively.

As an alternative application, we applied our platform to package digital information encoded in the DNA sequence for write-once, read-many archival DNA storage. Specifically, we cloned a sequence into the phPB52 variable domain that encoded a line from Act II, Scene 2 of <u>The Crucible</u> by Arthur Miller (phCruc; (40); **Figure 4b**). While the binary representation of this compressed and encrypted text file was converted into a DNA sequence using direct nucleotide conversion (A or G representing 0 and T or C representing 1), other encryption or compression approaches could alternatively be employed. A universal forward primer together with header information were added to the 5′ of the sequence, and an end-of-file (EOF), random slack space, and a universal reverse primer were added to the 3′ end (**Figure 4b**). The sequence was optimized for single-strandedness by ensuring no regions of sequence had greater than 8 bases that were repeated or complementary to any other region of the sequence. The DNA “memory block” was cloned into the phPB52 sequence (**Figure 4b**) and four-milliliter production of the phCruc phage showed 95% purity of the cssDNA as judged by agarose gel band intensities. Sanger sequencing was used to retrieve the insert sequence and decode the digital message (**External Table S1**).

## Discussion

We applied the *E. coli* str. M13cp strain for scalable bioproduction of pure cssDNA, which has the capabilities of generating isogenic material for biotechnological applications including scaffolded DNA origami and digital information archival and amplification, amongst other uses. The method employed here to direct purification of phage cssDNA without additional dsDNA contamination allows for new technology development in synthetic cssDNA sequence production that can be made bio-orthogonal and scalable, enabling future application to novel therapeutics and materials. Additional advances in the scaffolded DNA origami field are applicable to this strain, including the incorporation of DNAzymes (26) that would allow for greater control over the sequence and size of the produced linear ssDNA. However, in the approach used here, maintenance of the f1 origin and the selection marker in the produced phage allows for reinfection across the culture, which is important for subsequent biological amplification such as needed in phage display and, here, archival information storage. Moreover, circularization blocks exonuclease activity, which may prove important for therapeutic applications (41). In the future, improved understanding of phage biology should enable new approaches to excising specific coding sequences from M13 to generate engineered systems specifically designed for production of cssDNA.

The yields from the bioreactor approach used here were somewhat lower than wild type phage production that has been extensively optimized (26,27). This is due in part to the loss of the native feedback control over gene expression in the phage genome (42), the use of batch fermentation as opposed to a fed-batch approach that would allow for higher cell density (27), and plasmid loss due to ampicillin selection. Interestingly, we were not able to obtain clones of kanamycin or chloramphenicol selection cistrons on the vector purely under the control of the f1 origin. This may be due to the use of a single-stranded promoter, which might be overcome by alternative single-strand-specific promoters (43). This resistance insertion would then allow for fed-batch scale-up, leading to significantly improved yields.

Increased cssDNA production yields, together with advances in custom sequence design (25,26) and bio-orthogonality (44,45) with and without protein coding sequences, suggest that our approach is amenable to therapeutic applications in which ssDNA are used in circular (46) or linear (47) forms. In particular, scalable production of pure ssDNA at lower costs could enable yields required for therapeutic dosages of kilobase-length HDR template strands (48), a strategy that is further enabled by applying a DNAzyme approach for linearization (26). Scalable biological production of scaffolded DNA origami now matches production amounts from solid-state DNA synthesis commonly used for staple production, so that scaffolded DNA origami nanoparticles may now be produced at reasonable cost for mouse and higher animal therapeutic studies. Staple sequences synthesized with modifications to improve staple stability may further enhance nanoparticle lifetimes (28).

The alternative application of custom length and sequence scaffolds to encode digital information offers a write-once, read-many approach to low-cost massive archival data storage. Phage packaging is known to improve DNA stability against nuclease and chemical degradation (49), and ease of amplification in bacterial cultures makes this a intriguing method for native archival storage and biological-based information amplification, compatible with all sequencing strategies developed for M13 shotgun sequencing. Further, knowledge of phage biology and phage display offers an interesting set of possibilities for conditional amplification of phage sequences that encode specific digital information, a possible alternative to the current PCR-based solutions that are being developed.

## Materials and Methods

### Plasmid assembly by single-stranded DNA

All sequences of phage genomes (**External Table S1**) and primers (**External Table S2**) are contained in the External Excel document. The sequence of the f1 origin of replication was ordered from Integrated DNA Technologies (IDT, Inc., Coralville, IA) as a gBlock™ with 20 nt primers flanking the 5′ and 3′ sides designed to have a calculated melting temperature of 57°C (50). Double stranded DNA was generated by amplification of the synthetic gBlock f1 sequence with Phusion™ polymerase (New England Biolabs, Inc., Ipswitch, MA). The beta-lactamase (*bla*) ampicillin resistance gene with its promoter and terminator sequences were amplified from pUC19 using Phusion™ polymerase and 5′ and 3′ primers extended on their 5′ by the complementary pair of the f1 gBlock fragment. In each case, the PCR-amplified material was purified by ZymoClean agarose gel purification (Zymo Research, Inc., Irvine, CA) and column cleanup (Qiagen miniprep spin purification kit, Qiagen, Inc., Germany). Single-stranded DNA was generated using asymmetric production with 200 ng of purified dsDNA and 1 μM 5′-phosphorylated primer and Accustart HiFi polymerase (QuantaBio, Inc., Beverly, MA) in 1× Accustart HiFi buffer with 2 mM MgCl2, and cycled 25 times, as previously described (51). The ssDNA was gel- and column-purified. The two ssDNA products were then mixed in a 1:1 molar ratio and the ssDNA was converted to dsDNA using Phusion polymerase, column purified, and ligated using T4 DNA ligase (NEB) in 1× T4 DNA ligation buffer with 30 ng of amplified DNA incubated at room temperature overnight.

*E. coli* strains M13cp (33), DH5α F′Iq (Thermo Fisher, Inc., Waltham, MA), and SS320 (Lucigen, UK) were each made competent by washing log-phase grown cells in ice cold 100 mM CaCl_2_. 20 μL of competent cells were transformed with 2 μL of phagemid DNA ligation mix. Cells were incubated on ice for 30 minutes, heat shocked at 42°C for 45 seconds, and then put on ice. Pre-warmed SOB media was added and the cell culture was shaken at 37°C for 1 hour. 100 μL of cells were plated evenly across a Luria-Agar (LA) media plate made with 100 μg/mL ampicillin and 15 μg/mL chloramphenicol.

Individual colonies were selected and grown in 5 mL of Terrific Broth (TB) supplemented with 1% glycerol for overnight at 37°C. Bacteria was removed by centrifuging at 4,000 rpm for 10 minutes. Supernatant was removed and placed in a new 1.5 mL spin column and spun at 4,000 rpm for an additional 10 minutes. 1 μL of the clarified supernatant was added to 20 μL of nuclease-free water and heated to 95°C for 5 minutes, after which 1 μL of the heated solution was added to a Phusion PCR mix containing enzyme, buffer, nucleotides, and forward and reverse primers used to generate the plasmid. Positive colonies were determined by the presence of the PCR amplicon as visualized by agarose gel, and the purified phage were sent for Sanger sequencing. Of the eight colonies chosen, two were shown to have the correct sequence. The bacterial pellet was processed to purify all containing DNA by alkaline lysis and column purification (Qiagen miniprep spin kit, Qiagen, Inc., Germany).

Purified dsDNA was PCR amplified from phPB52 to have an EcoRI and PstI nuclease sites between the *bla* resistance cistron and the f1 ori. The PCR product was purified and digested alkaline phosphatase treated and gel purified. Synthetic DNA insert encoding digital information was generated using a computational algorithm that optimizes single-stranded compatibility. The full digital-DNA sequence and a partial sequence were amplified from a pUC19 vector containing the sequence to have flanking EcoRI and PstI sites, and digested with EcoRI and PstI nucleases. The product was gel purified. The inserts were ligated to the phPB52 digested vector in 1× T4 DNA ligation buffer (NEB) with 30 ng of vector DNA with three molar excess of synthetic inserts, incubated at room temperature overnight, and transformed into competent helper strain *E. coli*. These generated phPB84 and phCruc phage genomes.

### Synthetic phage production

Phage producing colonies, as judged by positive PCR, gel visualization, and sequencing results, were grown overnight in 4 mL 2×YT supplemented with 100 μg/mL ampicillin, 15 μg/mL of chloramphenicol and 5 μg/mL of tetracycline (Sigma-Aldrich, Inc.) in 15 mL culture tubes shaken at 200 RPMs at 37°C. The following day, the cultures were diluted to an O.D.600 of 0.05 in 2×YT supplemented with 100 μg/mL ampicillin, 15 μg/mL of chloramphenicol and 5 μg/mL of tetracycline and grown between 3 h to 27 h for time course experiments, and 8 h for media, pH, and strain optimization experiments. For pH optimization, the pH was controlled by addition of 100 mM HEPES-NaOH to the 2×YT media. Strain-specific antibiotics were used as recommended by the manufacture. After the chosen time point, the cultures were spun down at 4,000 RPMs for 15 minutes, after which the supernatant was removed to a fresh tube and spun at 4,000 RPMs for an additional 15 minutes, and filtered using a 0.45 μm cellulose acetate filter. For gel and nanodrop quantification, 400 μL of the clarified media was lysed by addition of Qiagen Buffer P1 supplemented with Proteinase K (20 μg/mL final; Sigma) and RNase A/T1 and incubated at 37°C for 1 h, followed by addition of Qiagen Buffer P2 and heating to 70°C for 15 minutes, and letting return to room temperature. Qiagen Buffer N3 was then added and precipitant was centrifuged. One volume of 100% ethanol was added to the supernatant and applied to a Qiagen spin column, and purified. The purified eluate DNA concentration was determined by A280 absorbance from a NanoDrop 2000 (Thermo Fisher) for each time point and condition tested, and ran on a 1% agarose gel in 1×Tris-Acetate-EDTA (TAE) stained with SybrSafe (Thermo Fisher) for visualization of the product. The ssDNA purity was judged by ImageJ (52) intensity analysis and the amount of ssDNA from the time point or condition was adjusted by this purity multiplied by the total amount of DNA found from A280 absorbance.

For milligram-scale production of synthetic miniphage, a Stedium Sartorius fermenter was used for growing 5 L of culture. An overnight culture was grown in 2×YT supplemented with 100 μg/mL Ampicillin and 15 μg/mL of chloramphenicol and 5 μg/mL of tetracycline and diluted to O.D. 600 of 0.05 for inoculating 5 L of media. The growth media for the batch fermentation was also 2×YT supplemented with 100 μg/mL Ampicillin and 15 μg/mL of chloramphenicol and 5 μg/mL of tetracycline. Oxygen and pH were monitored throughout the growth, and the pH was automatically adjusted with phosphoric acid and ammonium hydroxide, with a constant agitation of 800 RPM. Time points were taken approximately every hour and samples were processed as above for the shaker flask. At 8 h, 900 mL of liquid culture was removed for processing. For milligram-scale purification of ssDNA, 900 mL of liquid culture bacteria was pelleted by centrifuging twice at 4,000 × g for 20 min, followed by 0.45 μm cellulose acetate filtration. Phage from clarified media were precipitated by adding 6% w/v of polyethylene glycol-8000 (PEG-8000) and 3% w/v of NaCl and stirring continuously at 4°C for 1 h. Precipitated phage were collected by centrifuging at 12,000 × g for 1 h, and the PEG-8000 supernatant was removed completely, and pellet was resuspended in 30 mL of 10 mM Tris-HCl pH 8, 1 mM Ethylenediaminetetraacetic acid (EDTA) buffer (TE buffer). The phage was then processed using an EndoFree Maxiprep (Qiagen, Germany) column-based purification, following the manufacturer’s protocol with two adjustments. First, proteinase K (20 μg/mL final) was added to EndoFree Buffer P1 and incubated at 37°C for 1 h before addition of EndoFree Buffer P2 and incubation at 70°C for 10 min. The lysed phage was returned to room temperature before proceeding. Second, after removal of endotoxins, 0.2 v/v of 100% ethanol was added to the clarified sample, before applying to the EndoFree Maxiprep column to increase ssDNA binding. All other steps remained the same, and the cssDNA was eluted in 1 mL of endotoxin-free TE buffer. The amount of collected DNA was judged by absorbance at A280, and the purity was judged by running on a 1% agarose gel in 1× TAE stained with ethidium bromide.

Endotoxin amounts were tested using the ToxinSensor chromogenic LAL endotoxin assay kit (GenScript, Piscataway, NJ) following the manufacturer’s protocol. The cssDNA phPB84 was diluted to 10 nM in endotoxin-free water, with absorbance read on an Evolution 220 UV/Vis spectrophotometer (Thermo Fisher). Stability from exonuclease I degradation was tested by incubating cssDNA phPB84 with exonuclease I in 1× exonuclease buffer (NEB) at 37°C for 30 min. The reaction was quenched by incubating the reaction at 80°C for 15 min, and was subsequently ran on a 1% agarose gel in 1× TAE stained with ethidium bromide.

### DNA origami assembly

DNA purified from phages phPB52 and phPB84 were used to fold a pentagonal bipyramid with edge length 52 base pairs and 84 base pairs, respectively. Staples for each object were generated from the automated scaffold routing and staple design software DAEDALUS (19). Staples were synthesized by IDT, and listed in **External Table S3** and **S4**. To fold the nanoparticles, 20 nM of bacterially-produced and purified scaffold was incubated with 20-molar excess of staples in 1×TAE buffer with 12 mM MgCl2. The objects were annealed over 13 hours from 95°C to 24°C as previously described (19), and the folded particles were run on 1% agarose gel in 1×TAE buffer with 12 mM MgCl2 with the respective cssDNA scaffolds for reference. The folded nanoparticles were purified using a 100 kDA MWCO spin concentrator (Amicon) for a total of five 5-fold buffer exchanges as purification for TEM.

### Transmission electron microscopy

The structured DNA pentagonal bipyramid with 52-base-pair and 84-base-pair edge length assembled using the phage-produced scaffold was visualized by transmission electron microscopy (TEM). 200 μL of folded reaction was purified from excess staples and buffer exchanged into 20 mM Tris-HCl pH 8.0 and 8 mM MgCl_2_ using a 100kDa MWCO spin concentrator (Amicon, Merck Millipore, Billerica, MA). The concentration was subsequently adjusted to 5 nM. Carbon film with copper grids (CF200H-CU; Electron Microscopy Sciences Inc., Hatfield, PA) were glow discharged and the sample was applied for 60 seconds. The sample was then blotted from the grid using Whatman 42 ashless paper, and the grid was placed on drop of freshly prepared 1% uranyl-formate with 5mM NaOH for 10 s (53). Remaining stain was wicked away using Whatman 42 paper and dried before imaging. The grid was imaged on a Technai FEI with a Gatan camera.

## Supporting information

External Supplementary Tables

## Acknowledgements

We are grateful to Dr. Andrew Bradbury at Los Alamos National Lab for sharing the M13cp *E. coli* helper strain and to Dr. Longkuan Xiang at the Biomanufacturing Education and Training Center at the Worcester Polytechnic Institute for discussion and implementation of bacterial fermentation. Funding from the Office of Naval Research (N00014-14-1-0609; N00014-16-1-2181; N00014-16-1-2953) and the National Institutes of Health (1-R21-EB026008-01 and R01-MH112694) are gratefully acknowledged.

## Author Contributions

T.R.S. and M.B. initiated the study; T.R.S., H.H., R.D., and M.B. designed and analyzed growth experiments, T.R.S. and E.W. collected structural characterization data, T.R.S., R.R.D., and M.B. wrote the manuscript; All authors read and edited the manuscript.

## References

1. Messing, J., Crea, R. and Seeburg, P.H. (1981) A system for shotgun DNA sequencing. Nucleic Acids Res, 9, 309–321.

2. Zoller, M.J. and Smith, M. (1982) Oligonucleotide-directed mutagenesis using M13-derived vectors: an efficient and general procedure for the production of point mutations in any fragment of DNA. Nucleic Acids Res, 10, 6487–6500.

3. Chen, F., Pruett-Miller, S.M., Huang, Y., Gjoka, M., Duda, K., Taunton, J., Collingwood, T.N., Frodin, M. and Davis, G.D. (2011) High-frequency genome editing using ssDNA oligonucleotides with zinc-finger nucleases. Nat Methods, 8, 753–755.

4. Church, G.M., Gao, Y. and Kosuri, S. (2012) Next-generation digital information storage in DNA. Science, 337, 1628.

5. Goldman, N., Bertone, P., Chen, S., Dessimoz, C., LeProust, E.M., Sipos, B. and Birney, E. (2013) Towards practical, high-capacity, low-maintenance information storage in synthesized DNA. Nature, 494, 77–80.

6. Dietz, H., Douglas, S.M. and Shih, W.M. (2009) Folding DNA into twisted and curved nanoscale shapes. Science, 325, 725–730.

7. Douglas, S.M., Dietz, H., Liedl, T., Hogberg, B., Graf, F. and Shih, W.M. (2009) Self-assembly of DNA into nanoscale three-dimensional shapes. Nature, 459, 414–418.

8. Rothemund, P.W. (2006) Folding DNA to create nanoscale shapes and patterns. Nature, 440, 297–302.

9. Sharma, J., Chhabra, R., Cheng, A., Brownell, J., Liu, Y. and Yan, H. (2009) Control of self-assembly of DNA tubules through integration of gold nanoparticles. Science, 323, 112–116.

10. Diagne, C.T., Brun, C., Gasparutto, D., Baillin, X. and Tiron, R. (2016) DNA Origami Mask for Sub-Ten-Nanometer Lithography. ACS Nano, 10, 6458–6463.

11. Surwade, S.P., Zhao, S. and Liu, H. (2011) Molecular lithography through DNA-mediated etching and masking of SiO2. J Am Chem Soc, 133, 11868–11871.

12. Dutta, P.K., Levenberg, S., Loskutov, A., Jun, D., Saer, R., Beatty, J.T., Lin, S., Liu, Y., Woodbury, N.W. and Yan, H. (2014) A DNA-Directed Light-Harvesting/Reaction Center System. Journal of the American Chemical Society, 136, 16618–16625.

13. Dutta, P.K., Varghese, R., Nangreave, J., Lin, S., Yan, H. and Liu, Y. (2011) DNA-Directed Artificial Light-Harvesting Antenna. Journal of the American Chemical Society, 133, 11985–11993.

14. Hemmig, E.A., Creatore, C., Wünsch, B., Hecker, L., Mair, P., Parker, M.A., Emmott, S., Tinnefeld, P., Keyser, U.F. and Chin, A.W. (2016) Programming Light-Harvesting Efficiency Using DNA Origami. Nano Letters, 16, 2369–2374.

15. Pan, K., Boulais, E., Yang, L. and Bathe, M. (2014) Structure-based model for light-harvesting properties of nucleic acid nanostructures. Nucleic Acids Res, 42, 2159–2170.

16. Sun, W., Boulais, E., Hakobyan, Y., Wang, W.L., Guan, A., Bathe, M. and Yin, P. (2014) Casting inorganic structures with DNA molds. Science, 346, 1258361.

17. Douglas, S.M., Bachelet, I. and Church, G.M. (2012) A logic-gated nanorobot for targeted transport of molecular payloads. Science, 335, 831–834.

18. Zhao, Y.X., Shaw, A., Zeng, X., Benson, E., Nystrom, A.M. and Hogberg, B. (2012) DNA origami delivery system for cancer therapy with tunable release properties. ACS Nano, 6, 8684–8691.

19. Veneziano, R., Ratanalert, S., Zhang, K., Zhang, F., Yan, H., Chiu, W. and Bathe, M. (2016) Designer nanoscale DNA assemblies programmed from the top down. Science, 352, 1534.

20. Benson, E., Mohammed, A., Gardell, J., Masich, S., Czeizler, E., Orponen, P. and Hogberg, B. (2015) DNA rendering of polyhedral meshes at the nanoscale. Nature, 523, 441–444.

21. Douglas, S.M., Marblestone, A.H., Teerapittayanon, S., Vazquez, A., Church, G.M. and Shih, W.M. (2009) Rapid prototyping of 3D DNA-origami shapes with caDNAno. Nucleic Acids Res, 37, 5001–5006.

22. Jun, H., Shepherd, T.R., Zhang, K., Bricker, W.P., Li, S., Chiu, W. and Bathe, M. (2018) Automated Sequence Design of 3D Polyhedral Wireframe DNA Origami with Honeycomb Edges. Submitted.

23. Jun, H., Zhang, F., Shepherd, T., Ratanalert, S., Qi, X., Yan, H. and Bathe, M. (2019) Autonomously designed free-form 2D DNA origami. Sci Adv, 5, eaav0655.

24. Brown, S., Majikes, J., Martinez, A., Giron, T.M., Fennell, H., Samano, E.C. and LaBean, T.H. (2015) An easy-to-prepare mini-scaffold for DNA origami. Nanoscale, 7, 16621–16624.

25. Nafisi, P.M., Aksel, T. and Douglas, S.M. Construction of a novel phagemid to produce custom DNA origami scaffolds. Synthetic Biology, 3.

26. Praetorius, F., Kick, B., Behler, K.L., Honemann, M.N., Weuster-Botz, D. and Dietz, H. (2017) Biotechnological mass production of DNA origami. Nature, 552, 84–87.

27. Kick, B., Praetorius, F., Dietz, H. and Weuster-Botz, D. (2015) Efficient Production of Single-Stranded Phage DNA as Scaffolds for DNA Origami. Nano Lett, 15, 4672–4676.

28. Conway, J.W., McLaughlin, C.K., Castor, K.J. and Sleiman, H. (2013) DNA nanostructure serum stability: greater than the sum of its parts. Chem Commun (Camb), 49, 1172–1174.

29. Vieira, J. and Messing, J. (1987) Production of single-stranded plasmid DNA. Methods Enzymol, 153, 3–11.

30. Pasqualini, R. and Ruoslahti, E. (1996) Organ targeting in vivo using phage display peptide libraries. Nature, 380, 364–366.

31. Winter, G., Griffiths, A.D., Hawkins, R.E. and Hoogenboom, H.R. (1994) Making antibodies by phage display technology. Annu Rev Immunol, 12, 433–455.

32. Ferrara, F., Kim, C.Y., Naranjo, L.A. and Bradbury, A.R. (2015) Large scale production of phage antibody libraries using a bioreactor. MAbs, 7, 26–31.

33. Chasteen, L., Ayriss, J., Pavlik, P. and Bradbury, A.R. (2006) Eliminating helper phage from phage display. Nucleic Acids Res, 34, e145.

34. Reddy, P. and McKenney, K. (1996) Improved method for the production of M13 phage and single-stranded DNA for DNA sequencing. Biotechniques, 20, 854–856, 858, 860.

35. Bryksin, A.V. and Matsumura, I. (2010) Overlap extension PCR cloning: a simple and reliable way to create recombinant plasmids. Biotechniques, 48, 463–465.

36. van den Ent, F. and Lowe, J. (2006) RF cloning: a restriction-free method for inserting target genes into plasmids. J Biochem Biophys Methods, 67, 67–74.

37. Veneziano, R., Shepherd, T.R., Ratanalert, S., Bellou, L., Tao, C. and Bathe, M. (2018) In vitro synthesis of gene-length single-stranded DNA. Sci Rep, 8, 6548.

38. Dotto, G.P., Horiuchi, K. and Zinder, N.D. (1984) The functional origin of bacteriophage f1 DNA replication. Its signals and domains. J Mol Biol, 172, 507–521.

39. Hahn, J., Wickham, S.F., Shih, W.M. and Perrault, S.D. (2014) Addressing the instability of DNA nanostructures in tissue culture. ACS Nano, 8, 8765–8775.

40. Miller, A. (1953). The Crucible. Act II, Scene 2.

41. Lovett, S.T. (2011) The DNA Exonucleases of Escherichia coli. EcoSal Plus, 4.

42. Smeal, S.W., Schmitt, M.A., Pereira, R.R., Prasad, A. and Fisk, J.D. (2017) Simulation of the M13 life cycle II: Investigation of the control mechanisms of M13 infection and establishment of the carrier state. Virology, 500, 275–284.

43. Masai, H. and Arai, K. (1997) Frpo: a novel single-stranded DNA promoter for transcription and for primer RNA synthesis of DNA replication. Cell, 89, 897–907.

44. Kozyra, J., Ceccarelli, A., Torelli, E., Lopiccolo, A., Gu, J.Y., Fellermann, H., Stimming, U. and Krasnogor, N. (2017) Designing Uniquely Addressable Bio-orthogonal Synthetic Scaffolds for DNA and RNA Origami. ACS Synth Biol, 6, 1140–1149.

45. Rovner, A.J., Haimovich, A.D., Katz, S.R., Li, Z., Grome, M.W., Gassaway, B.M., Amiram, M., Patel, J.R., Gallagher, R.R., Rinehart, J. et al. (2015) Recoded organisms engineered to depend on synthetic amino acids. Nature, 518, 89–93.

46. Seidl, C.I. and Ryan, K. (2011) Circular single-stranded synthetic DNA delivery vectors for microRNA. PLoS One, 6, e16925.

47. Davis, L. and Maizels, N. (2016) Two Distinct Pathways Support Gene Correction by Single-Stranded Donors at DNA Nicks. Cell Rep, 17, 1872–1881.

48. Richardson, C.D., Ray, G.J., DeWitt, M.A., Curie, G.L. and Corn, J.E. (2016) Enhancing homology-directed genome editing by catalytically active and inactive CRISPR-Cas9 using asymmetric donor DNA. Nat Biotechnol, 34, 339–344.

49. Clark, J.R. and March, J.B. (2006) Bacteriophages and biotechnology: vaccines, gene therapy and antibacterials. Trends Biotechnol, 24, 212–218.

50. SantaLucia, J.Jr., (1998) A unified view of polymer, dumbbell, and oligonucleotide DNA nearest-neighbor thermodynamics. Proc Natl Acad Sci U S A, 95, 1460–1465.

51. Veneziano, R., Shepherd, T.R., Bellou, L., Ratanalert, S., Tao, C. and Bathe, M. (2017) Enzymatic synthesis of gene-length single-stranded DNA. bioRxiv.

52. Schneider, C.A., Rasband, W.S. and Eliceiri, K.W. (2012) NIH Image to ImageJ: 25 years of image analysis. Nat Methods, 9, 671–675.

53. Castro, C.E., Kilchherr, F., Kim, D.-N., Shiao, E.L., Wauer, T., Wortmann, P., Bathe, M. and Dietz, H. (2011) A primer to scaffolded DNA origami. Nature Methods, 8, 221–229.

