## Supplementary Information for "Bioproduction of single-stranded DNA from isogenic miniphage"

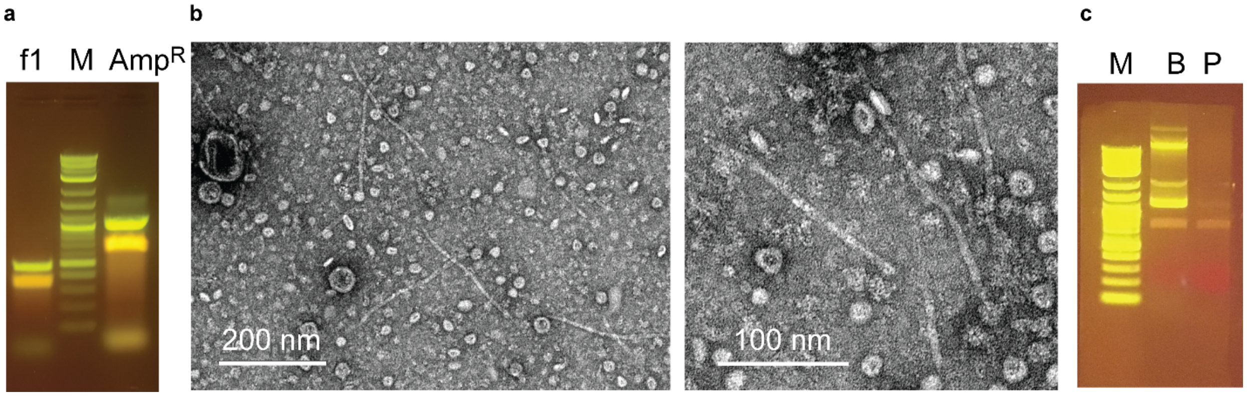


**Figure S1. Single-strand DNA synthesis and phage assembly.** (**a**) Full agarose gel from **Figure 1b**, showing asymmetric PCR products of f1 origin-containing synthetic gBlock and bla-selection marker-containing ssDNAs. (**b**) Wide-field TEM micrograph showing assembled phage in the clarified media, rod-shaped and approximately 200 nm in size. (**c**) Full agarose gel from **Figure 1d** showing DNA purification results from the bacterial pellet and the purified phage from the media. M: Marker. Scale bar is shown for reference.

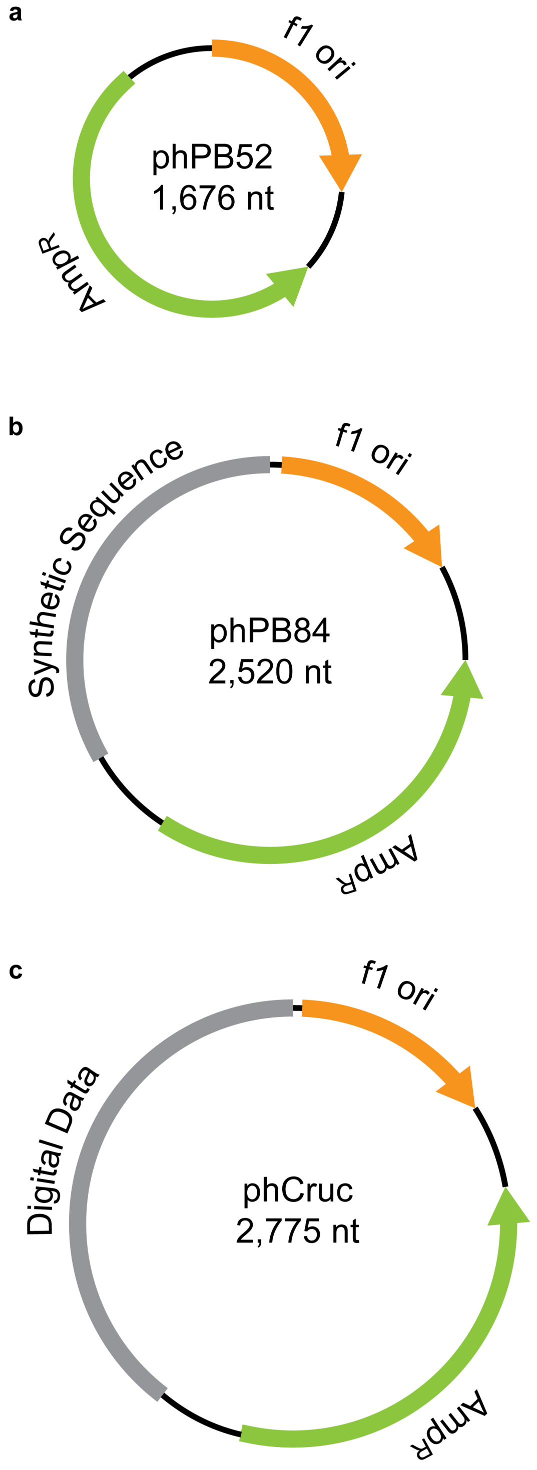


**Figure S2. Plasmid maps of constructs generated.** (**a**) phPB52, (**b**) phPB84, and (**c**) phCruc.

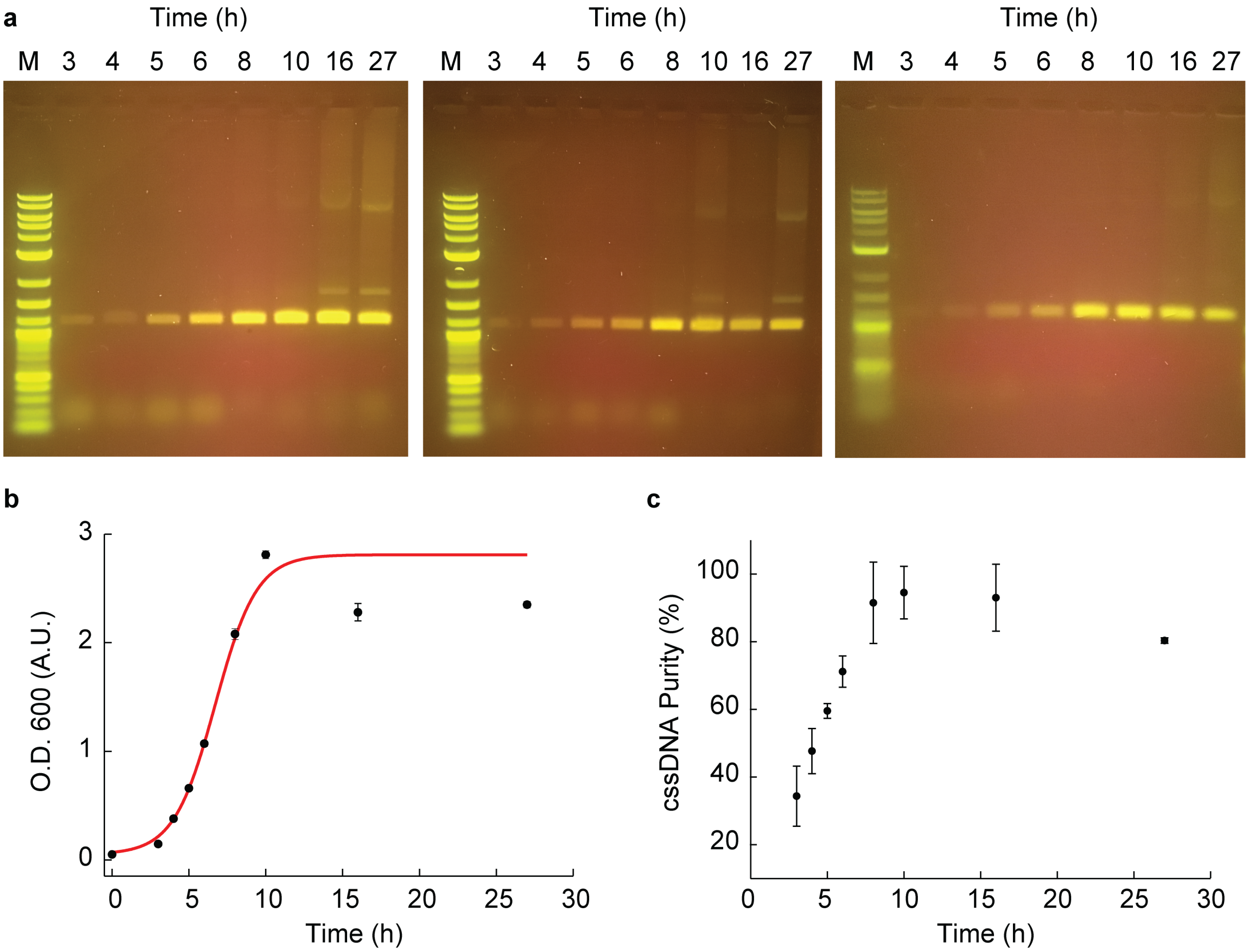


**Figure S3. Triplicate time course assays for optimizing phPB84 miniphage production in a shaker flask.** (**a**) DNA was prepared from column purification of processed, cleared supernatant of bacterial growths at the indicated times. Growth was in 2×YT media using the SS320 strain transformed with phPB84 and the M13cp helper plasmid. (**b**) O.D.600 time course monitoring the growth of the bacterial culture, inoculated at the 0 time point to have an O.D.600 of 0.05. (**c**) Percent purity was measured based on gel percent intensity of the cssDNA compared to total lane intensity, measured by ImageJ (52)

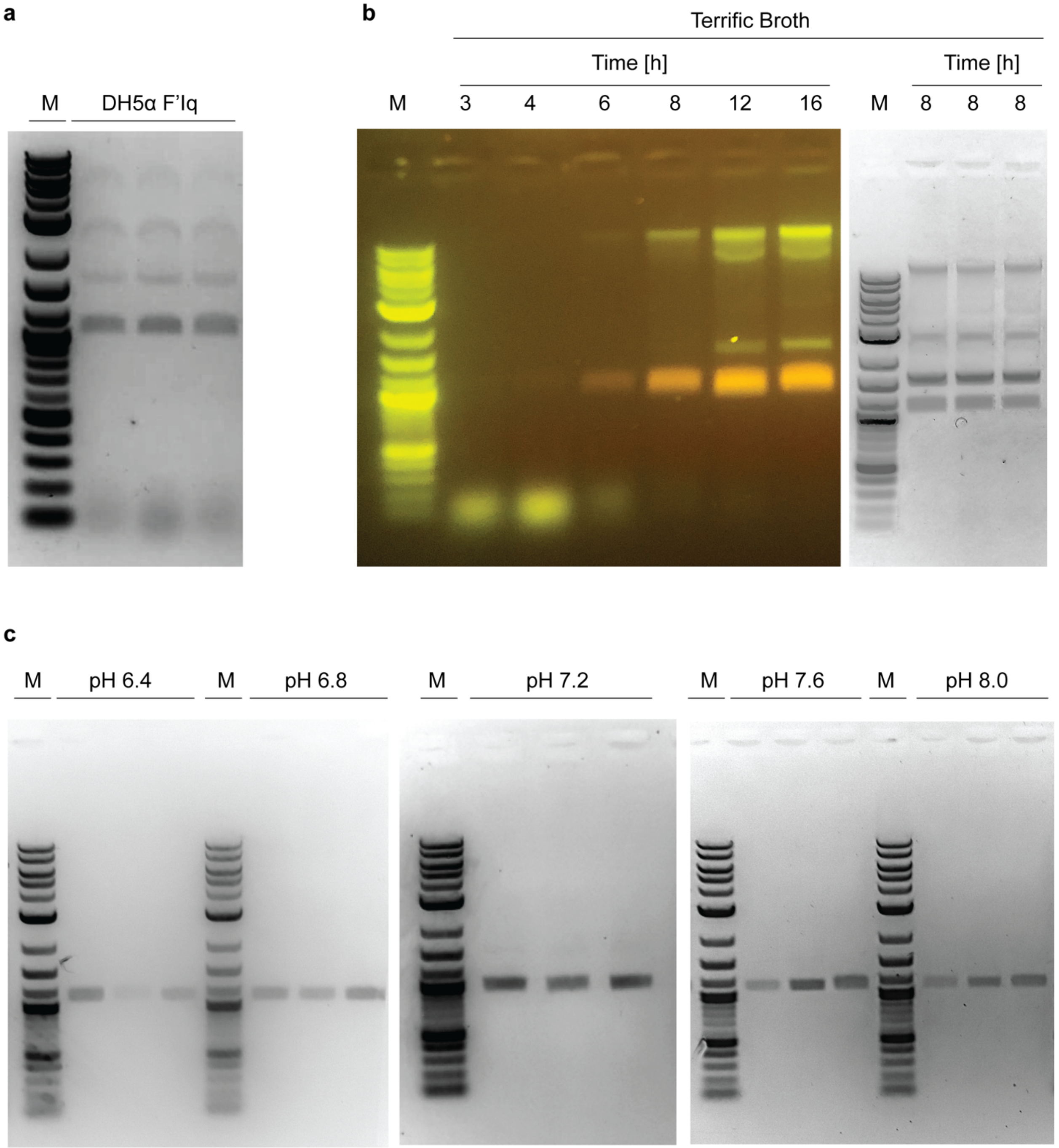


**Figure S4. Agarose gels showing triplicate measurements of cssDNA prepared from the cleared media for optimizing conditions.** (**a**) Independent colonies of the DH5a F'Iq (Invitrogen) strain with helper plasmid transformed with phPB84 and grown to the 8 h time point in 2×YT media. (**b**) Agarose gel showing processed cssDNA from a time course assay and triplicate growths at the 8 h time point with Terrific Broth as the growth media using the SS320 strain. (**c**) Agarose gel showing triplicate cssDNA prepared from the cleared media using 2×YT media at pHs indicated, buffered with 100 mM HEPES-NaOH at the 8 h time point using the SS320 strain.

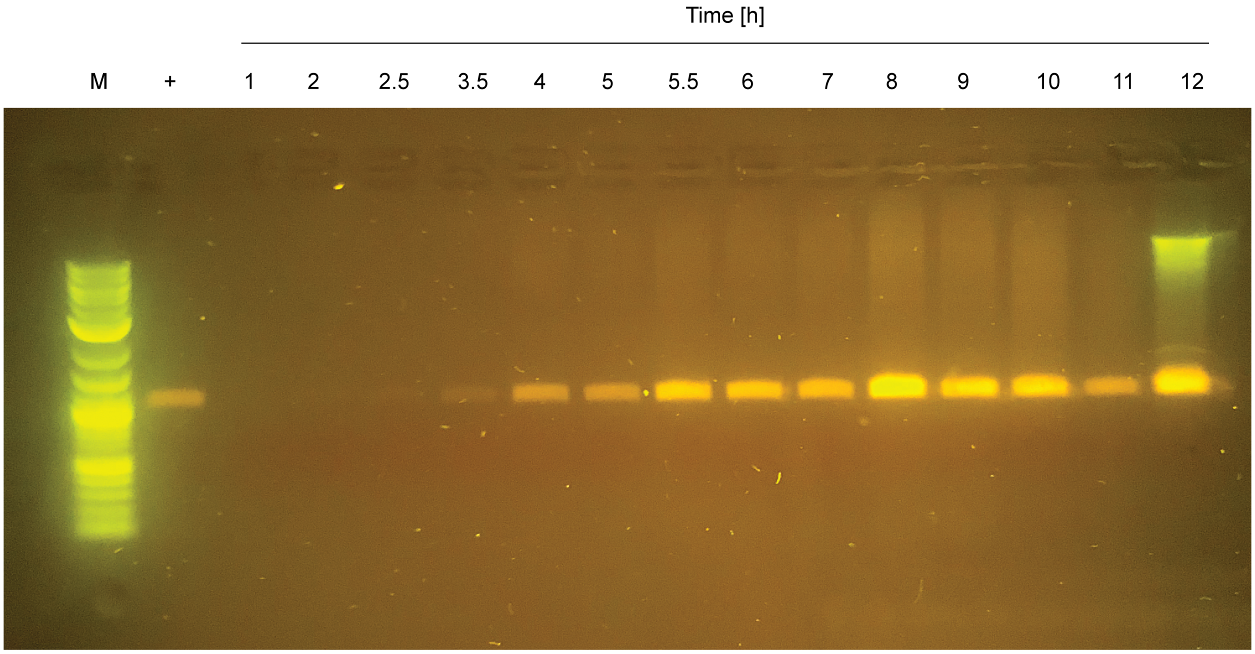


**Figure S5. Agarose gel of time course production of cssDNA phPB84 in a bioreactor.** Growth was in 2×YT media using the SS320 strain transformed with phPB84 and the M13cp helper plasmid with pH controlled to 7.0. M: marker; +: phPB84 from shaker flask growth.

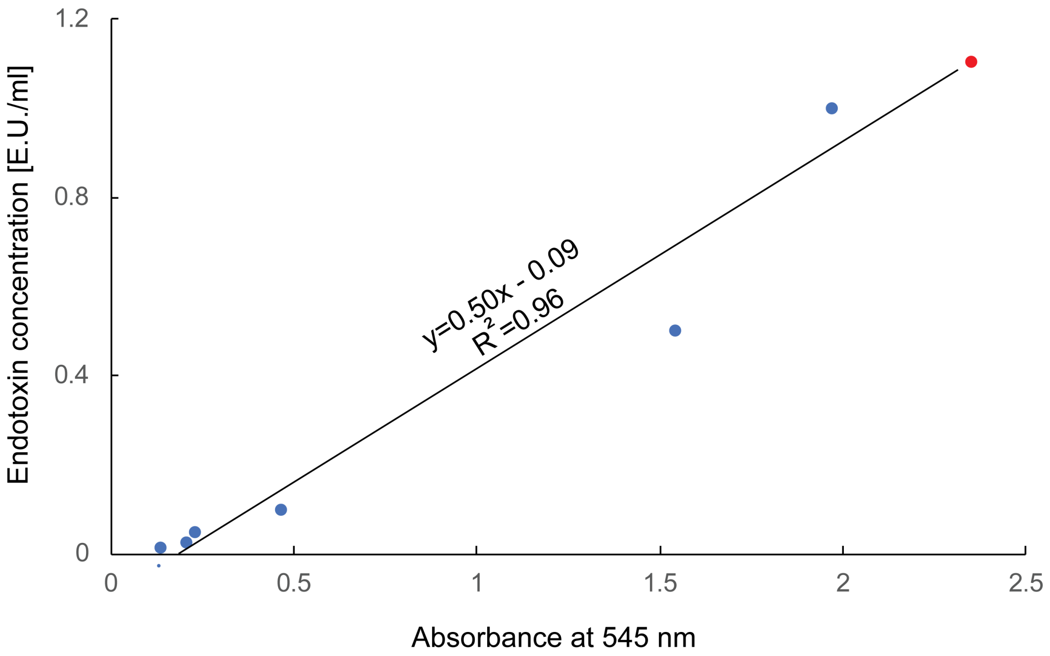


**Figure S6. Endotoxin screening using standard curve with the 10 nM scaffold data point shown.** The ToxinSensor Chromogenic LAL Endotoxin Assay Kit (GenScript) was used to assay endotoxin levels in the prepared DNA. The eluted DNA was diluted to 10 nM from the Endofree Maxiprep purification kit (Qiagen) and tested against a standard curve by following the manufacturer's protocol monitoring absorbance at 545 nm, shown as blue dots. The endotoxin level was calculated to be 1.1 ± 0.1, shown as a red dot.

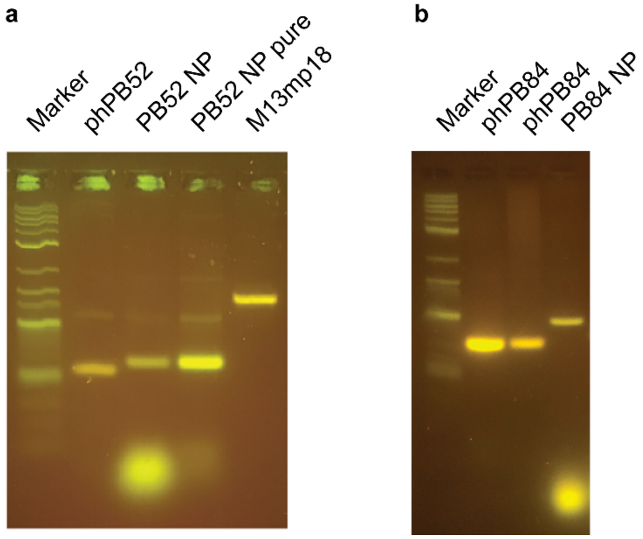


**Figure S7. Agarose gel analysis of folding of nanoparticles.** (**a**) Uncropped gel of a folded pentagonal bipyramid of 52-bp edge length. (**b**) Uncropped gel of a folded pentagonal bipyramid of 84-bp edge length.

a

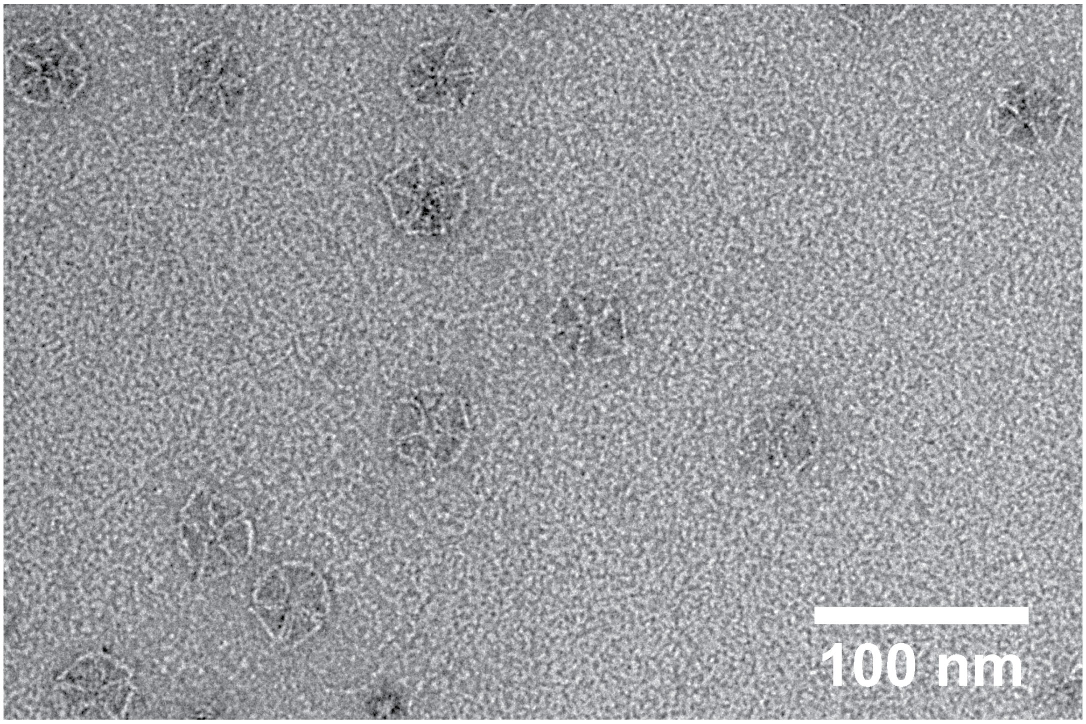


b

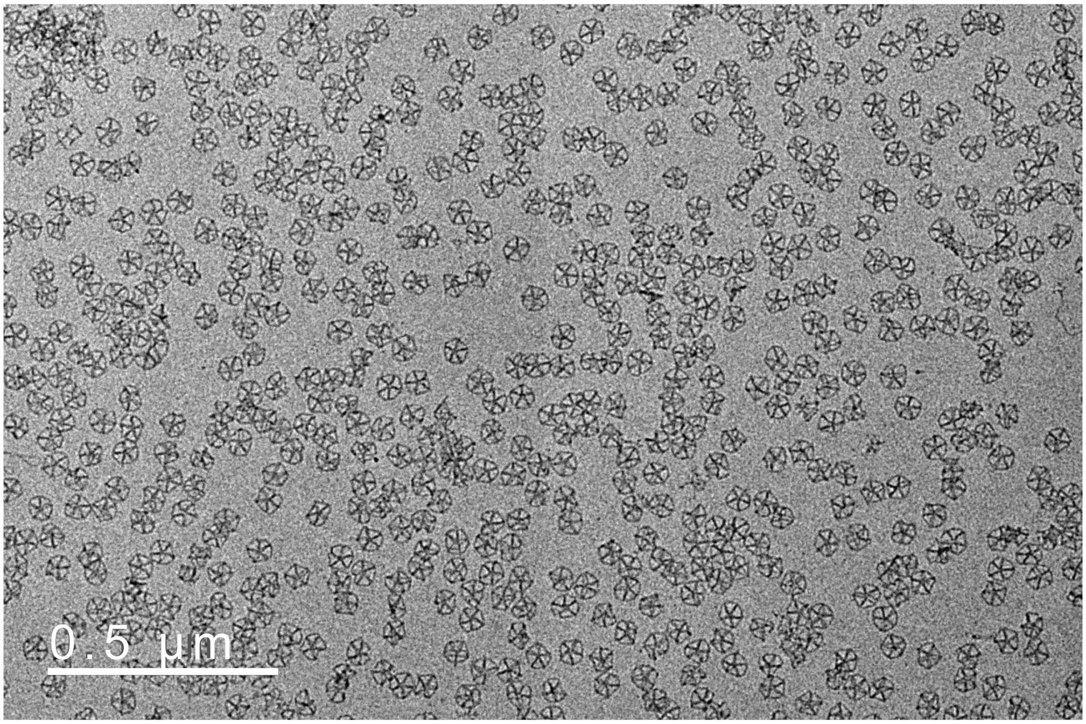


**Figure S8. Wide-field TEM micrographs of scaffolded DNA nanoparticles.** (**a**) Pentagonal bipyramid with 52-bp edge lengths and (**b**) Pentagonal bipyramid with 84-bp edge lengths are shown. Scale bars are shown for reference.

**External Table S1. Sequences of each of the phages and amplicons generated in this study.**

**External Table S2. Primers used for cloning and sequence validation.**

**External Table S3. Staples for 52-bp edge-length pentagonal bipyramid nanoparticle from phPB52 sequence**

**External Table S4. Staples for 84-bp edge-length pentagonal bipyramid nanoparticle from phPB84 sequence.**
